# Low rank mechanisms underlying flexible visual representations

**DOI:** 10.1101/730978

**Authors:** Douglas A. Ruff, Cheng Xue, Lily E. Kramer, Faisal Baqai, Marlene R. Cohen

## Abstract

Neuronal population responses to sensory stimuli are remarkably flexible. The responses of neurons in visual cortex depend on stimulus properties (e.g. contrast), processes that affect all stages of visual processing (e.g. adaptation), and cognitive processes (e.g attention or task switching). The effects of all of these processes on trial-averaged responses of individual neurons are well-described by divisive normalization, in which responses are scaled by the total stimulus drive. Normalization describes how a staggering variety of sensory, cognitive, and motor processes affect individual neurons (1), but whether different normalization processes could be mediated by the same mechanism remains poorly understood. We and others recently showed that attention has low rank effects on the covariability of populations of neurons in visual area V4 (2–4), which strongly constrains mechanistic models mechanism (2). We hypothesized that measuring changes in population covariability associated with other normalization processes could clarify whether they might share a mechanism. Our experimental design included measurements in multiple visual areas using four normalization processes. We found that contrast, adaptation, attention, and task switching affect the responses of populations of neurons in primate visual cortex in a similarly low rank way. These results suggest that a given circuit uses a common mechanism to perform many forms of normalization and likely reflect a general principle that applies to a wide range of brain areas and sensory, cognitive, or motor processes.

## Introduction

Understanding the biological basis of a neural computation can clarify the cognitive processes by which our brains convert information about the sensory world into action. Divisive normalization, in which the responses of individual neurons are divisively scaled by the mean drive to the population, is a simple computation that explains a wide variety of sensory, cognitive, and motor processes. In the visual system, normalization accounts for the modulation involving changes to the visual stimulus (e.g. stimulus contrast or surround suppression; (1, 5–12), modulation originating from the earliest stages of visual processing in the retina (e.g. adaptation; (1, 13–16), and modulation originating from cognitive processes internal to the nervous system (e.g. attention, task switching, learning, task difficulty, or multisensory integration (17–28).

The existence of normalization in multiple species, brain areas, and functional processes led to the appealing hypothesis that these processes share a common underlying mechanism (1). However, the normalization equation is not a mechanistic model, and the divisive scaling of trial-averaged responses is consistent with many models.

In contrast, neuronal population responses can be used to constrain models. We showed recently that measuring how trial-to-trial response variability is shared across a neuronal population and how that covariability depends on sensory or cognitive processes provides strong constraints on a model of visual cortex. We and others have shown that covariability of firing rates in visual cortex is typically low rank (2–4, 29–35). This means that shared variability is well-described as a low-dimensional process that affects neurons with different weights rather than higher order interactions between neurons or subpopulations. Furthermore, we showed that attention has an even lower rank effect on covariability (approximately rank one), as evidenced by the observation that the relationship between noise and signal correlation is largely unchanged by attention (36) and by the results of a variety of methods for directly measuring the rank attention-related modulation of shared variability (2–4).

Unlike in the data, many typical models of cortical circuits (including balanced excitatory-inhibitory networks with fast inhibition or with slow inhibition with broad connectivity) produce high rank variability (2, 37, 38). We and our collaborators showed recently that requiring the models to have realistic timescales and connectivity and constraining parameters by oberved effects of attention on covariability places strong constraints on the underlying mechanisms (2, 39).

We therefore hypothesized that measuring how different normalization-associated processes affect population covariability may reveal whether they could plausibly be mediated by the same mechanism. We designed our experiment to simultaneously record the effects of multiple normalization-associated processes on the same neuronal population and also to evaluate the generality of our findings by recording from multiple brain areas using different behavioral tasks. We therefore have two data sets. Our first data set consists of simultaneously recorded effects of contrast, adaptation, and spatial attention on the responses of small populations of neurons in visual area V4. We chose those three processes for three reasons. First, their responses on the trial-averaged responses of V4 neurons are well-described by divisive normalization (1). Second, they are all known to affect the extent to which trial-to-trial response variability is shared between pairs of visual neurons (36, 40–44), which makes it possible to compare their effects on the dimensionality of correlated variability. Third, and most importantly, these three modulatory processes represent a strong test of the hypothesis that all processes involving normalization involve a common mechanism because they originate at different stages of visual processing (contrast is a change in the visual stimulus and affects neuronal responses at all processing stages; adaptation affects responses beginning in the retina, and endogenous attention more strongly affects later stages of visual processing; Figure 1A). We found that although the way each of contrast, adaptation, and attention modulate a given neuron’s mean response was uncorrelated with modulation by other factors, all three of these processes affect covariability in a low rank way.

**Figure 1.**
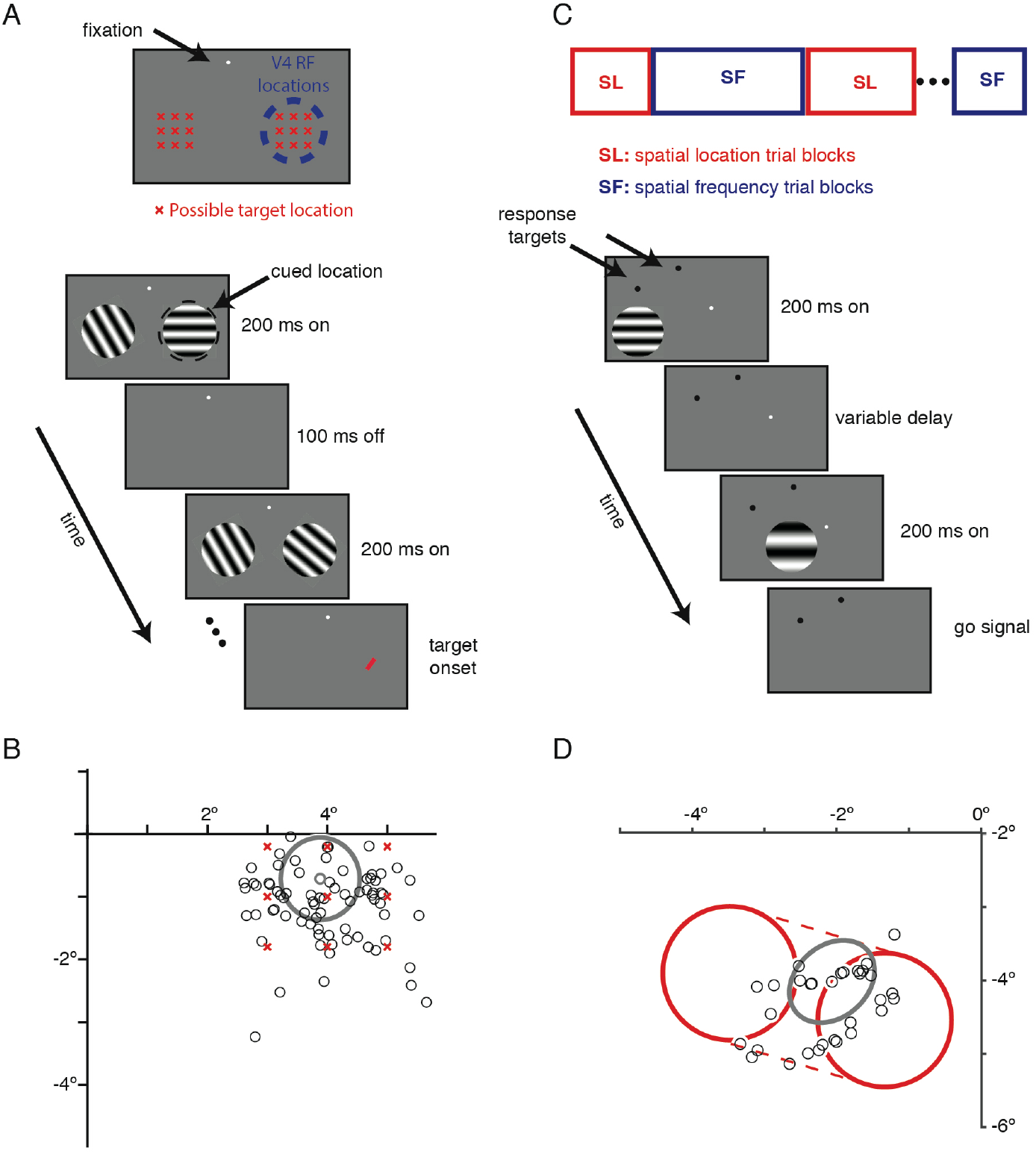
Methods. A. Layout of possible target locations in the cued bar detection task and schematic of the detection task. Possible locations are marked with red Xs in top panel. The animal maintained fixation while static Gabor stimuli flashed on and off in various configurations. The animal was rewarded for directing a saccade to the target stimulus (red bar) within 450ms of its appearance. B) Estimates of receptive field centers from an example V4 recording session (black circles) during the bar detection task. An estimate of the size of one example unit’s receptive field is drawn in gray. Possible target locations are depicted by red Xs. C) Schematic of the task-switching task. two subsequent briefly displayed moving Gabors differ both in their spatial locations and spatial frequencies, only one of which is behaviorally relevant in each trial. The monkey is rewarded, in different trial blocks, to discriminate the change in the relevant feature while ignoring the change in the irrelevant feature. D) Estimates of receptive field centers from an example V1 recording session (black circles) during the task-switching paradigm. An estimate of the size of one example unit’s receptive field is shown by a gray ellipse. The stimulus size and range of stimulus locations are shown in red.

Our second data set shows that these low rank effects were not limited to area V4 or to processes that have homogeneous effects on large populations: we also found that even when different visual features of a stimulus are encoded by the same group of primary visual cortical (V1) neurons, changing their behavioral relevance produces low rank effects on neuronal response variability. Together, these data are consistent with the idea that despite the differences between contrast, adaptation, attention, and task switching, they affect the responses of populations of neurons in primate visual cortex in similar ways. More broadly, they suggest that simultaneous recordings from groups of neurons will be key for understanding the biological bases of a large range of neuronal computations.

## Results

### Simultaneous psychophysical and neurophysiological measurements of contrast, adaptation, and attention

To simultaneously measure the effects of contrast, adaptation, and attention on neuronal populations, we trained two monkeys (Macaca mulatta) to perform the cued detection task illustrated in Figure 1. A trial began when the animal fixated a central spot. While the animal waited to detect the onset of a target (see Methods), we presented a series of pairs of static gratings across a range of orientations (200 ms duration per stimulus, one stimulus per hemifield; Figure 1A). The animal was rewarded for making a saccade to a small red bar that could appear at one of 9 possible locations in each hemifield within 450 ms of its onset or for maintaining fixation if no bar was presented after 8000 ms. The gratings varied in orientation and contrast. The animal was cued in blocks of trials using unanalyzed instruction trials as to which hemifield the bar was likely to appear. The attention cue accurately predicted the side that the bar would appear on subsequent trials 85% of the time.

The animals’ task performance shows that they respected the attention cue. The animals correctly detected a greater proportion of target bar stimuli when they were presented at one of the nine locations in the cued (attended) hemifield than one of the nine locations in the uncued hemifield. Overall, the animals detected 84% of target bars in the cued hemifield and 78% in the uncued hemifield (t-test, p<0.05).

While the monkeys performed the detection task, we measured the effects of contrast, adaptation, and attention on neuronal populations in visual area V4 using chronically implanted microelectrode arrays (see Methods; Figure 1B). We recorded single and multiunit activity during daily experimental sessions for several weeks in each animal while the animals performed the detection task. The most important aspect of our design was to measure contrast, adaptation, and attention simultaneously by sorting the same data in different ways. We measured contrast by comparing responses to stimuli with high and low contrast, adaptation by comparing responses to the first and second presentation of a given orientation, and attention by comparing responses when attention was directed toward or away from the grating in the same hemifield as the units’ receptive fields.

### Task switching: a framework for generalizing our results to different brain areas and tasks

While contrast, adaptation, and adaptation are maximally different in terms of the stage of visual processing in which they originate, they have something important in common in the context of our experiment. All three processes involve a nonspecific drive to the populations of neurons we recorded. We reasoned that if we were ever going to see higher rank changes in population response variability, it might be in a situation in which the modulatory process had qualitatively different effects on different subsets of neurons.

To test the generality of our results, we recorded the responses of neurons in a different visual area (primary visual cortex, or V1) while two different monkeys performed a two-interval, forced choice task in which they had to switch which of two features (the spatial location or the spatial frequency of the Gabor patch, both of which are encoded in V1) they discriminated. In alternating blocks of trials, they indicated whether the spatial frequency of the Gabor stimuli increased or decreased or whether the location of the stimuli moved to the left or right (Figure 1C and 1D). The change amounts were chosen so that the difficulties of the two tasks would be comparable (monkey 3 average performance: 75% for spatial location task, 69% for spatial frequency task; monkey 4 average performance: 79% for spatial location task, 80% for spatial frequency task; no significant pairwise performance differences for the two tasks across sessions: p=0.47 for monkey 3, p=0.37 for monkey 4, Wilcoxon signed rank test). The neurons we recorded from encode information about both of the two features, presumably in separable encoding dimensions. Therefore, by comparing the covariability in trial blocks where different encoding dimensions are behaviorally relevant, we can test if higher rank changes are necessary for more complicated modulations.

### Contrast, adaptation, and attention have diverse and largely separable effects on individual units and pairs of units

Consistent with previous results, we found that contrast (8, 10, 45, 46), adaptation (13, 15, 47), and spatial attention (48–50) were all associated with changes in the mean firing rates of the V4 units we recorded from. On average, contrast and attention increased mean rates (Figure 2A; mean physiologists’ index comparing high and low contrast = 0.17 and mean index comparing attention toward and away from the hemifield containing the unit’s receptive field for attention = 0.01, two-tailed Wilcoxon signed rank test, p= effectively 0 and p=2.3×10^−12^, respectively), and adaptation decreased mean rates (mean index comparing the first and second presentation of the same orientation = 0.01, two-tailed Wilcoxon signed rank test, p=3.3×10^−11^).

**Figure 2.**
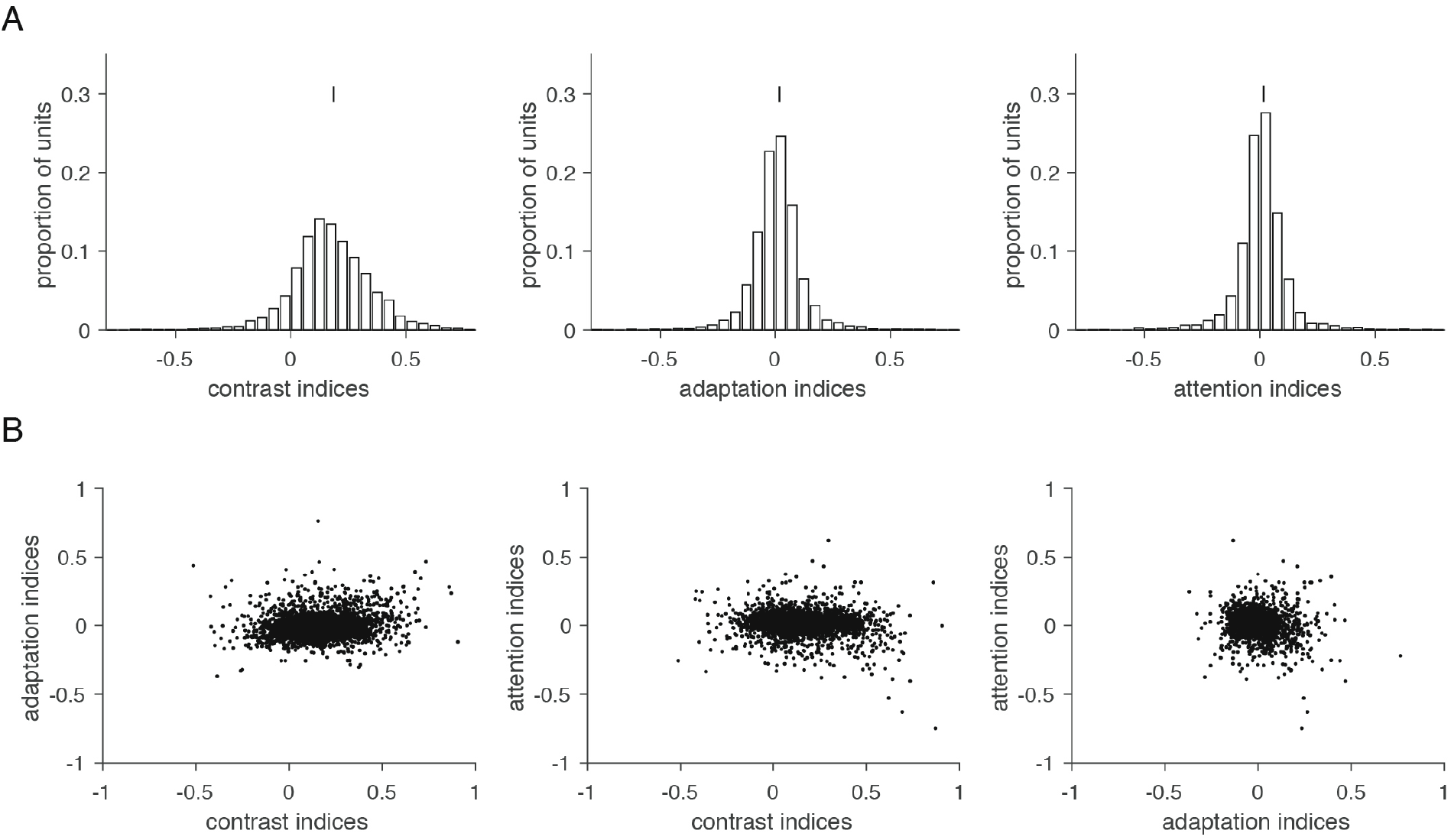
Contrast, adaptation and attention affect firing rates in a largely separable manner. A) physiologist indices (difference divided by the sum of responses in the two conditions) for each unit for contrast, adaptation, and attention. Distribution means indicated by vertical line. 5,328 units from 10 sessions. B) The effects of contrast, adaptation and attention on individual units are weakly related.

Our data suggest that contrast, adaptation, and attention affect the firing rates of individual units in a largely separable way (17, 51–59). The extent to which each unit’s mean rate was modulated by one of the three processes was weakly, but significantly correlated with modulation by the other processes (Figure 2B; Pearson’s correlation coefficients between contrast and adaptation indices = 0.13, between contrast and attention indices = −0.1, and between adaptation and attention = −0.12, p=7.2×10^−20^, p=4.4×10^−12^ and p=9.2×10^−19^, respectively). These results suggest that contrast, adaptation, and attention are associated with robust changes to mean rates, but do not provide strong evidence that these changes equally target the same subsets of neurons.

Previous studies have demonstrated that a variety of processes modulate the strength of covariance or pairwise correlations (often termed r_SC_; (60) in sensory areas. These include anesthesia (34, 61, 62), attention (36, 44, 63), arousal, alertness or task difficulty (64–66), and locomotion (67). Consistent with these reports, we found that contrast, adaptation, and attention were associated with changes in covariability (36, 40–44); Figure 3A, changes in covariability associated with all three modulatory processes were significantly different from 0; Wilcoxon signed rank test, p<0.01).

**Figure 3.**
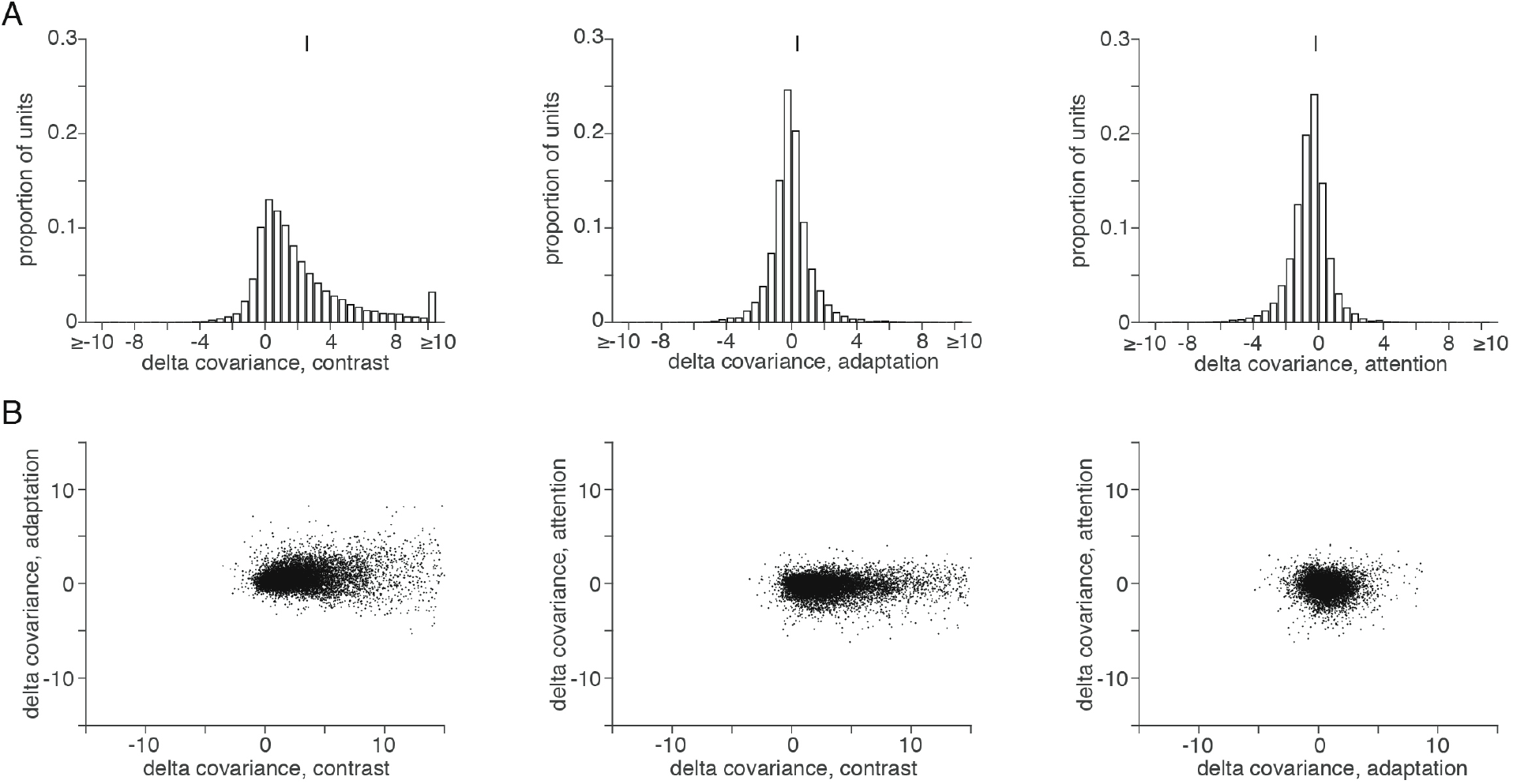
Contrast, adaptation and attention affect correlated variability between pairs of units. A. Histograms of covariance changes for contrast, adaptation, and attention. Distribution means indicated by vertical line. 20,774 pairs from 10 sessions. B) Covariance changes associated with contrast, adaptation and attention on individual units are weakly related.

On average, contrast and adaptation increased covariance (Figure 3A; mean physiologists’ index comparing high and low contrast = 2.43 and mean index comparing the first and second presentation of the same orientation = 0.19, two-tailed Wilcoxon signed rank test p=0 and p=5.9×10^−66^, respectively), and attention decreased covariance (mean index comparing attention toward and away from the hemifield containing the unit’s receptive field for attention = −0.30 which is significantly less than 0, two-tailed Wilcoxon signed rank test p=1.4×10^−163^). The effects of attention, adaptation, and contrast on covariance were not strongly interrelated (Figure 3C; the Pearson’s correlation between each unit’s mean change in covariance with all simultaneously recorded units = −0.07 between adaptation and attention, 0.001 between contrast and attention, and 0.22 between contrast and adaptation, two-sided t-test, p=1.3×10^−13^, p=0.29, and p=3.9×10^−128^, respectively).

### Contrast, adaptation, and attention are all associated with low rank modulation of response covariability

We reasoned that if contrast, adaptation, and attention are associated with the same computation, they might all affect covariability in similarly low rank ways and along similar dimensions of neuronal population activity. We used factor analysis on the z-scored responses to each stimulus to evaluate the rank of the changes in response variability associated with each modulatory process. The majority of the population variance is contained in the first few modes (the majority of the variance is concentrated in the first eigenmode; Figure 4A). As in previous studies (2–4), we found that attention primarily affects variability in the first eigenmode (Figure 4A, compare black and red lines). Consistent with the idea that modulatory processes affect the same small number of dimensions, we also found that contrast and adaptation affected variability predominantly in this first mode (Figure 4A, compare black and blue lines and black and green lines, respectively).

**Figure 4.**
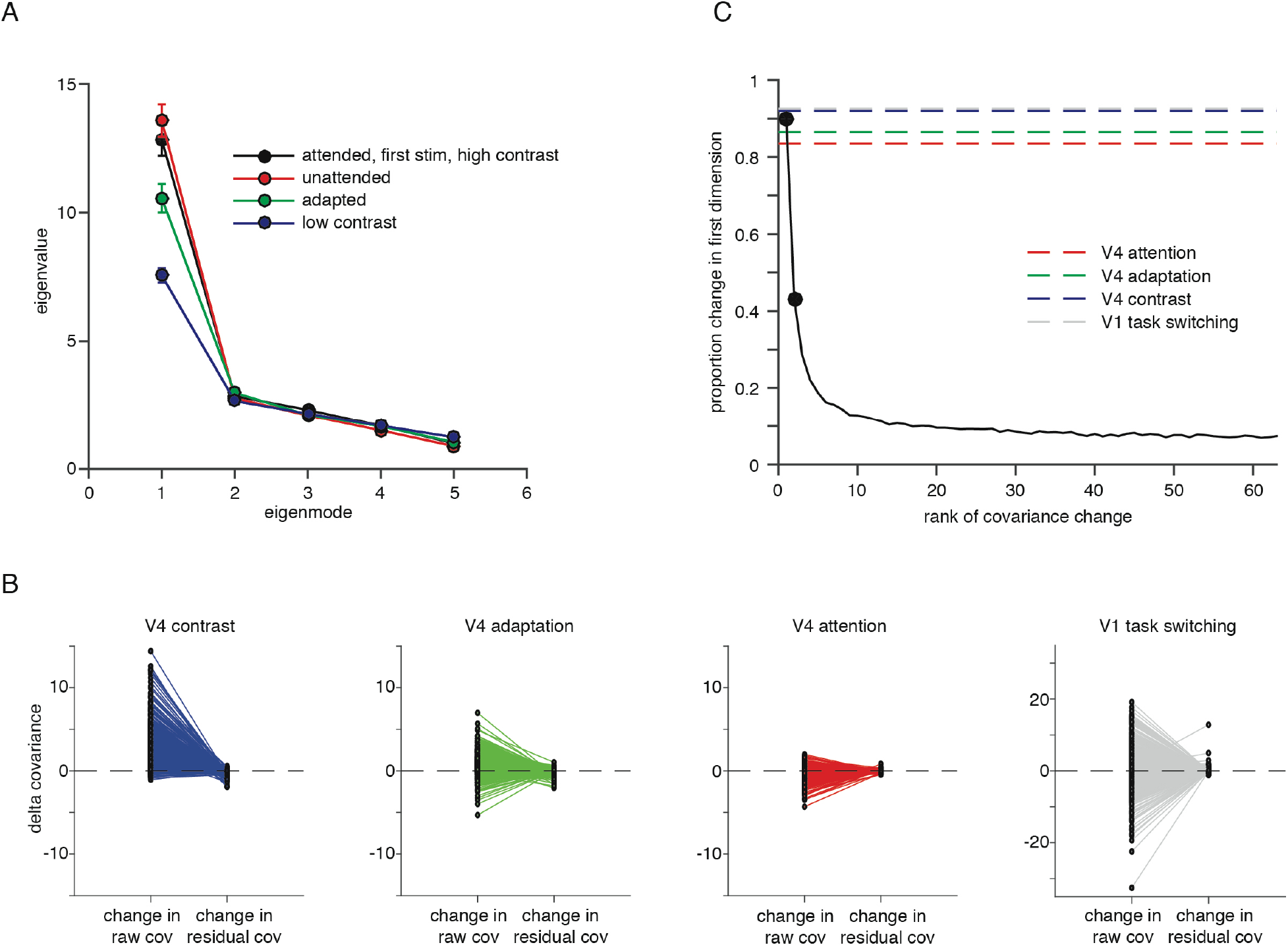
Contrast, adaptation and attention affect population responses in a small number of dimensions. A) eigenvalues (shared variance) of the population covariance matrix corresponding to the first five modes calculated using factor analysis (33, 68). B) Changes in average covariance associated with contrast, adaptation and attention in V4 or task switching in V1 between each unit’s raw (left) or residual (first eigenmode removed) responses and all other simultaneously recorded units. Positive numbers mean that covariance was higher for high contrast, the first stimulus, the attended stimulus, or the location discrimination task, respectively. C) Proportion change in covariance in the first eigenmode as a function of the rank of the covariance change in a simulation. We constructed simulated data sets using the mean number of units and trials as our real data sets. We constructed changes in covariance of different ranks using a procedure described in Methods. We then analyzed the simulated data the same way as the real data in Figure 4B and plotted 1-the ratio of the covariance change in the residual and raw data. The mean values for each of the data sets in Figure 4B and for the V1 data are plotted as horizontal lines in the corresponding color.

These changes in covariability associated with contrast, adaptation, and attention can be completely explained by changes in a single dimension of population response space. The changes in pairwise covariance associated with each process are essentially abolished when the first eigenmode is removed (2); Figure 4B, mean proportion of total change in covariance for V4 = 0.84 (attention), 0.87 (adaptation), and 0.92 (contrast)). The magnitude of this change is consistent with a change in a single dimension, given the size of our data set (Figure 4C). In principle, if we recorded from much larger populations, we might find modulations that span more than a single dimensions. However, these analyses show that while contrast, adaptation, and attention are associated with diverse (and different signed) modulations of mean responses and pairwise covariability in visual cortex, they all modulate responses in the same, small number of dimensions. These results are therefore consistent with the hypothesis that all processes associated with normalization affect populations of V4 neurons via similar mechanisms.

### Low rank modulation of response variability is not limited to homogeneous processes

One possibility is that the modulation we observed is low rank because attention, adaptation, and contrast have relatively homogeneous effects on the groups of neurons we recorded. For example, contrast typically increases the responses of V4 neurons, making it possible that contrast has a low rank, monolithic effect on population responses. A related possibility is that trial-to-trial variability in these processes (such as, for example, the animal’s internal attention state) is the source of this low rank variability (4, 69). The different sign relationships between the strength of the low rank modulation and performance (e.g. good performance is associated with low variability for attention but high variability for adaptation and contrast) make it unlikely that the explanation is that simple, but measuring trial-to-trial variability in internal states is notoriously difficult.

We conducted a strong test of the hypothesis that even complex cognitive processes are associated with low rank modulation by measuring modulation of visual cortical activity associated with switching between two tasks that are mediated by the same population of neurons (Figure 1C). Switching between spatial frequency and location discrimination tasks caused both increases and decreases in firing rates (54.2% of units had higher mean rates in the spatial frequency than spatial location task) and pairwise covariance (57.0% of units had higher mean r_SC_ in the spatial frequency than spatial location task).

We found that task switching affected the covariability of groups of V1 neurons in a low rank manner that was very analogous to the effects of attention, adaptation, and contrast on groups of V4 neurons (Figure 4B). Like in V4, the covariance change was almost exclusively oriented along the primary axis (mean proportion of total change in covariance for V1 (task switching) = 0.92). These results suggest that the low rank modulation of population covariability we observed in V4 is not limited to extrastriate cortex or to processes with relatively homogeneous effects on neuronal populations.

## Discussion

### Constraints on mechanistic models

We showed that although attention, adaptation, contrast, and task switching have quantitatively and qualitatively different effects on the responses of visual cortical neurons, they all affect shared response variability in a low rank way. This observation is important because it places strong constraints on models of how modulatory processes affect cortical circuits.

We and our collaborators showed (2) that although many models produce variability whose magnitude matches observed pairwise noise correlations, the only model that produces low rank fluctuations is one in which inhibition is slower than excitation (which is physiologically realistic; 70, 71–73) and in which the connectivity of inhibitory neurons is spatially restricted (which is also realistic; 74, 75, 76). In that model, attention (and presumably contrast and adaptation, although these were not modeled explicitly) could exert low rank changes in shared variability through an input whose effect is to change the balance between excitation and inhibition, increasing the activity or influence of inhibitory neurons relative to excitatory ones (2). This simple mechanism is a good candidate for a general mechanism mediating a broad class of normalization computations in primate visual cortex.

The idea that all normalization processes utilize the same mechanism appears to be contradicted by two recent results showing that in mouse visual cortex, normalization primarily involves changes in excitation (77), but in the Drosophila antenna lobe, normalization is mediated through changes in inhibition (78). Our model leaves room for both of these possibilities: normalization is accomplished by shifting the E/I balance, which could involve changes to excitation, inhibition, or both (2, 39). One possibility is that both systems use the same broad mechanism (changing E/I balance) but employ different means to change that balance. Another possibility is that each *circuit* (e.g. primate V1 or V4) uses the same mechanism for all forms of normalization, but that different systems use different mechanisms.

A recent set of studies suggested a mechanism in which each neuron has a set of inputs that perform normalization and that attention and other modulatory processes act through those inputs (20–24, 79, 80). This idea is supported by the observation that the extent to which the mean responses of individual neurons in visual cortex are modulated by switching attention between two stimuli within their receptive fields is highly correlated with the magnitude of response suppression associated with placing an additional stimulus in the receptive field (20–22, 24, 79). However, this strong correlation does not seem to apply to all modulatory processes: those same authors showed that modulation by feature attention is not correlated with multi-stimulus suppression (27), and we showed that modulation by contrast, adaptation, and attention are uncorrelated.

Together, these observations suggest that while all forms of normalization may involve the same or similar circuit mechanisms, the involvement of particular neurons likely depends on the specific modulatory process or circuit. In some form, the idea that the involvement of specific neurons depends on the specific context is necessary to explain the observations that the extent of modulation by, for example, feature attention, depends on how closely a neurons’ tuning matches the attended stimulus(81–85).

### Implications for information coding

A curious consequence of the observation that modulatory processes affect response variability in a low rank way is that those modulations should have little to no effect on the stimulus information that could be read out of a neuronal population by an optimal decoder. An elegant series of theoretical studies showed that shared variability affects the Fisher information encoded in a neuronal population only when it is aligned with the dimensions in population space that are being read out, and that in large populations, the amount of shared variability that is aligned with those dimensions is vanishingly small (86–88). If contrast, adaptation, and attention affect only a small number of dimensions in population space, their effects could easily be discounted by an optimal decoder.

Why then are attention, adaptation, contrast, and task switching associated with such large changes in visual perception? Our results are consistent with only two possibilities: 1) that the modulations in visual cortex associated with these processes do not affect perception, or 2) that readout is suboptimal in a particular way, such that the first mode (which is associated with the changes in shared variability) plays a larger role than it would in an optimal readout scheme. While the first possibility has been proposed for attention (in favor of an alternative model in which attention is mediated through changes in the visual information that is communicated to downstream areas involved in decision-making; (89–92), it seems implausible for a low level process like contrast. Contrast is associated with changes from the earliest stages of visual processing, which are then communicated to visual cortex. The idea that contrast-related changes in visual cortex are not associated with the concomitant changes in perception seems implausible.

On its surface, the second possibility, that visual information is read out suboptimally such that low rank changes in variability affect perception, seems equally implausible. Any readout mechanism approaching optimality should have access to a much higher dimensional representation of visual information, which would minimize the impact of low rank changes in response variability. However, we showed recently that in an orientation change detection task, monkeys’ choices were much more closely aligned with the axes in V4 population space that correspond to shared variability than would be expected if decoding approached optimality (93). One possibility is that, either because of a biological constraint or a need to optimize something other than performance on a specific task (89), responses oriented along the same dimensions that affect modulatory processes carry an outsized influence on perception. Determining the validity of these two possibilities will be an important avenue for further work.

## Methods

### Subjects and recording methods

The subjects in our V4 experiments were two adult male rhesus monkeys (*Macaca mulatta*, 11 and 9 kilograms). The subjects in our V1 experiments were two different adult male rhesus monkeys (*Macaca mulatta*, 10 and 12 kilograms). All animal procedures were approved by the Institutional Animal Care and Use Committees of the University of Pittsburgh and Carnegie Mellon University. Before training, we implanted each animal with a titanium head post. Monkey 1 from the V4 experiment had previously been trained to perform a direction change detection task (93) and had a 10×10 microelectrode array (Blackrock Microsystems) implanted in area V4 in one cerebral hemisphere 4 months prior to the start of training for the current study. After completion of the previous study, the monkey was retrained for 4 weeks to perform the bar detection task. Monkey 2 for the V4 experiment was trained for 4 months to perform the bar detection task before we implanted a 10×10 microelectrode array and began recordings. Both V1 monkeys were trained to perform the two discrimination tasks (Figure 1C) before we implanted a 6×8 array in area V1. In all cases, the distance between adjacent electrodes was 400 μm, and each electrode was 1 mm long. We identified areas V1 or V4 using stereotactic coordinates and by visually inspection of sulcal landmarks. Each array was connected to a percutaneous connector that allowed electrophysiological recordings.

To simultaneously measure the effects of contrast, adaptation, and attention on neuronal populations, two monkeys performed the detection task illustrated in Figure 1A. A trial began when the animal fixated a central spot. The animal maintained fixation while waiting to detect the onset of a small red bar (100 ms stimulus presentation, the bar was .2° thick and ranged from .25° to .5° long, with pixel intensities selected at the start of each session to make the task sufficiently difficult) presented after a delay period picked from an exponential distribution (minimum time before target onset was 300 ms, tau ranged from 3800 - 4500 ms between sessions, max trial time was 8000 ms). While waiting for the target, a series of pairs of static gratings (200 ms duration, 100 ms interflash interval) were presented at different orientations; one in the joint receptive field of the recorded neurons and the other in the opposite hemifield. We manipulated contrast by changing the contrast of the gratings (we used full contrast and low contrast stimuli which ranged between 5% and 20% across sessions), adaptation by comparing gratings that were presented following a grating of the same or different orientation, and attention by cueing the animal in blocks of trials as to which hemifield to expect the red target bar to appear (the cue was valid on 85% of trials). The animal was rewarded for making a saccade to the red bar within 450 ms of its onset or for maintaining fixation if no bar was presented after 8000 ms.

To measure the effects of task switching on populations of V1 neurons, we trained two additional monkeys to perform the discrimination tasks illustrated in Figure 1C. The animals maintained fixation on a central spot when two subsequent moving Gabor patches with randomly drawn spatial locations and spatial frequencies were displayed for 200ms each, separated by a variable delay (100~150ms for monkey 3, 300~500ms for monkey 4). Trials were grouped in alternating blocks of two behavioral contexts (each block contains 225 trials for monkey 3, and 40 trials on average for monkey 4). In the spatial location context, the monkeys discriminated the difference in the two stimuli’s locations (left or right shift), while disregarding their spatial frequencies; likewise in the spatial frequency context, the monkeys discriminated the difference in the two stimuli’s spatial frequencies (increase or decrease), while disregarding their spatial locations. The monkeys were trained to make a saccade to corresponding saccade targets to report their discrimination decision.

We presented visual stimuli on a calibrated CRT monitor (calibrated to linearize intensity, 1,024 × 768 pixels, 120-Hz refresh rate) placed 57 cm from the animal. We monitored eye position using an infrared eye tracker (Eyelink 1000, SR Research). We used custom software (written in Matlab using the Psychophysics Toolbox, ref) to present stimuli and monitor behavior. We recorded eye position and pupil diameter (1,000 samples per s), neuronal responses (30,000 samples per s) and the signal from a photodiode to align neuronal responses to stimulus presentation times (10,000 samples per s) using hardware from Ripple.

To measure the effects of modulatory processes on neuronal populations, we recorded single and multiunit activity from Utah arrays during daily experimental sessions for several weeks in each animal (V4 experiments: 7 sessions in Monkey 1 and 11 sessions in Monkey 2; V1 experiments: 21 sessions in Monkey 3 and 24 sessions in Monkey 4). Using our recording methods, it is nearly impossible to tell whether we recorded from the same single- or multi-unit clusters on subsequent days and with the exception of summary figures 2 and 3, our analyses focus on within session questions. We spike sorted single units and multi-unit clusters manually following the experiment using Plexon's Offline Sorter and we combined single and multiunits for all analyses (we use the term “units” to refer to either). For the recordings in V4, we positioned the gratings and possible target bar locations such that they fell in the joint receptive fields of most units (Figure 1B). We included V4 experimental sessions for neuronal analysis if we recorded from a minimum of ten units simultaneously whose average visual responses to the first stimulus shown (which was not used for any subsequent analyses) on all completed high contrast trials was at least 25% more than baseline and also at least 5 spikes per second greater than baseline (7 sessions from Monkey 1 and 2 sessions from Monkey 2; average of 63 responsive units per session and 29 stimulus repetitions of each orientation and condition).

For the analysis of the V4 data, we counted spikes in response to visual stimuli in 200ms bin shifted 50ms to account for the visual latencies of V4. Our analyses included all responses after the initial stimulus presentation (to remove contrast adaptation; 36) from all trials and were not baseline subtracted. We included responses from all completed trials up until 250ms before a behavioral event (i.e., the target onset or a fixation break). For each unit (for measures of individual unit rates) or pair of units recorded simultaneously but not from the same electrode (for covariance), we computed physiologists’ indices (defined as the difference between responses in two conditions divided by their sum), covariance, or r_SC_ separately for each unique combination of grating orientation, contrast, adaptation, and attention condition and averaged the results as described in the figure legends. For the indices in Figure 2A, we compared activity using the responses from the attend out, low contrast, adapted condition as the comparison point against the responses in the condition where one of those features was changed, for contrast, adaptation and attention, respectively. For the scatter plots in Figure 2B, average indices for each unit were calculated for all pairs of stimulus conditions and then collapsed across conditions. Covariances in Figure 3 were calculated with a similar principle as in Figures 2A and average covariance changes for each pair of units were calculated within all pairs of stimulus conditions (as in Figure 3A) and then collapsed across conditions.

For recordings in V1, the positions of the stimuli are chosen so that they overlap with most of the receptive fields of V1 neurons (Figure 1D). We included V1 experimental sessions for neuronal analyses if there are at least 50 trials with identical initial stimuli (same location and spatial frequency) for each condition. All V1 units were included in the analyses as long as their stimulus responses were significantly different from baseline activity. (13 sessions from Monkey 3 and 20 sessions from Monkey 4; average of 53 responsive units per session and 153 repetitions of identical initial stimuli for each condition per session. We included all completed trials (where monkeys successfully maintained fixation until they made a saccade and indicated their choice). We analyzed the response to the first stimulus display only (i.e. the number of spikes within the 200ms of stimulus time window shifted 34 ms for the visual latencies of V1 (94).

### Population analyses

We used factor analysis to identify the dimensions in population space that account for the most shared variance in the population. For the V4 data, we began by z-scoring the spike count responses of each neuron to gratings of each orientation in each contrast, adaptation, and attention condition. For the V1 data, we z-scored the responses to the each unique stimulus, using responses to the first stimulus on each trial (to avoid adaptation effects). We then combined the data across all conditions to form a number of neurons x number of trials z-scored response matrix which we call X. We used Factor analysis (68, 95) to identify the five dimensions in population space that accounted for the greatest variability in population responses. Because we z-scored the responses, these dimensions represent the dimensions of greatest covariability in the population. We computed the residual covariance (Figures 4B) after subtracting the first mode as Cov(X,X)-L1xL1’ as in (2), where X is the population response matrix, Cov(X,X) is the raw covariance matrix, and L1 is the loading matrix when fitting with a single factor. We repeated this analysis on multivariate Gaussian distributions with mean covariance equal to the means for the two contrast conditions, which was the manipulation associated with the largest covariance change in our study (Figure 4C).

We conducted simulations to evaluate the proportion of the covariance change in the first dimension that would be measured in our data sets with *n* units if the true covariance change was of rank *k*. We constructed a covariance matrix *S*=*W*W’+D*, where *W* is a random matrix of size *k*×*n* and *D* is a random diagonal matrix with positive elements. We then drew the responses of each simulated neuron on each trial from a multivariate Gaussian distribution with covariance *S* and analyzed the results as we did our real data.

## References

1. Carandini M & Heeger DJ (2012) Normalization as a canonical neural computation. Nature Reviews Neuroscience 13(1):51–62.

2. Huang C, et al. (2019) Circuit Models of Low-Dimensional Shared Variability in Cortical Networks. Neuron 101(2):337–348.

3. Rabinowitz NC, Goris RL, Cohen M, & Simoncelli EP (2015) Attention stabilizes the shared gain of V4 populations. Elife 4:e08998.

4. Ecker AS, Denfield GH, Bethge M, & Tolias AS (2016) On the Structure of Neuronal Population Activity under Fluctuations in Attentional State. J Neurosci 36(5):1775–1789.

5. Ayaz A, Saleem AB, Scholvinck ML, & Carandini M (2013) Locomotion controls spatial integration in mouse visual cortex. Curr Biol 23(10):890–894.

6. Nassi JJ, Gomez-Laberge C, Kreiman G, & Born RT (2014) Corticocortical feedback increases the spatial extent of normalization. Front Syst Neurosci 8:105.

7. Carandini M & Heeger D (1994) Summation and division by neurons in primate visual cortex. Science 264(5163):1333–1336.

8. Carandini M, Heeger DJ, & Movshon JA (1997) Linearity and normalization in simple cells of the macaque primary visual cortex. The Journal of neuroscience 17(21):8621–8644.

9. Simoncelli EP & Schwartz O (1999) Modeling surround suppression in V1 neurons with a statistically-derived normalization model. Advances in Neural Information Processing Systems, eds Kearns MS, Solla SA, & Cohn DA (MIT Press, Cambridge, MA), Vol 11, pp 153–159.

10. Heeger DJ (1992) Normalization of cell responses in cat striate cortex. Visual Neuroscience 9(2):181–197.

11. Busse L, Wade AR, & Carandini M (2009) Representation of concurrent stimuli by population activity in visual cortex. Neuron 64(6):931–942.

12. Britten KH & Heuer HW (1999) Spatial summation in the receptive fields of MT neurons. The Journal of Neuroscience 19(12):5074–5084.

13. Solomon SG & Kohn A (2014) Moving sensory adaptation beyond suppressive effects in single neurons. Curr Biol 24(20):R1012–1022.

14. Shapley R & Enroth-Cugell C (1984) Visual adaptation and retinal gain controls. Prog Ret Res 3:263–346.

15. Kohn A (2007) Visual adaptation: physiology, mechanisms, and functional benefits. J Neurophysiol 97(5):3155–3164.

16. Olveczky BP, Baccus SA, & Meister M (2007) Retinal adaptation to object motion. Neuron 56(4):689–700.

17. Sanayei M, Herrero JL, Distler C, & Thiele A (2015) Attention and normalization circuits in macaque V1. European Journal of Neuroscience 41(7):949–964.

18. Ohshiro T, Angelaki DE, & DeAngelis GC (2011) A normalization model of multisensory integration. Nature neuroscience 14(6):775–782.

19. Ohshiro T, Angelaki DE, & DeAngelis GC (2017) A Neural Signature of Divisive Normalization at the Level of Multisensory Integration in Primate Cortex. Neuron 95(2):399–411 e398.

20. Lee J & Maunsell JHR (2009) A normalization model of attentional modulation of single unit responses. PloS one 4(2):e4651.

21. Ni AM & Maunsell JHR (2017) Spatially tuned normalization explains attention modulation variance within neurons. J Neurophysiol 118(3):1903–1913.

22. Reynolds JH & Heeger DJ (2009) The normalization model of attention. Neuron 61(2):168–185.

23. Ruff DA & Cohen MR (2017) A normalization model suggests that attention changes the weighting of inputs between visual areas. Proc Natl Acad Sci U S A 114(20):E4085–E4094.

24. Ni AM, Ray S, & Maunsell JHR (2012) Tuned normalization explains the size of attention modulations. Neuron 73(4):803–813.

25. Louie K, Khaw MW, & Glimcher PW (2013) Normalization is a general neural mechanism for context-dependent decision making. Proceedings of the National Academy of Sciences of the United States of America 110(15):6139–6144.

26. Yamada H, Louie K, Tymula A, & Glimcher PW (2018) Free choice shapes normalized value signals in medial orbitofrontal cortex. Nat Commun 9(1):162.

27. Ni AM & Maunsell JHR (2019) Neuronal effects of spatial and feature attention differ due to normalization. J Neurosci.

28. Boynton G (2009) A framework for describing the effects of attention on visual responses. Vision research 49(10):1129–1143.

29. Goris RLT, Movshon JA, & Simoncelli EP (2014) Partitioning neuronal variability. Nature neuroscience 17(6):858–865.

30. Lin IC, Okun M, Carandini M, & Harris KD (2015) The Nature of Shared Cortical Variability. Neuron 87(3):644–656.

31. Cowley BR, Smith MA, Kohn A, & Yu BM (2016) Stimulus-Driven Population Activity Patterns in Macaque Primary Visual Cortex. PLoS Comput Biol 12(12):e1005185.

32. Semedo JD, Zandvakili A, Machens CK, Yu BM, & Kohn A (2019) Cortical Areas Interact through a Communication Subspace. Neuron.

33. Williamson RC, et al. (2016) Scaling Properties of Dimensionality Reduction for Neural Populations and Network Models. PLoS Comput Biol 12(12):e1005141.

34. Ecker AS, et al. (2014) State dependence of noise correlations in macaque primary visual cortex. Neuron 82(1):235–248.

35. Stringer C, et al. (2016) Inhibitory control of correlated intrinsic variability in cortical networks. Elife 5.

36. Cohen MR & Maunsell JHR (2009) Attention improves performance primarily by reducing interneuronal correlations. Nature Neuroscience 12(12):1594–1600.

37. Shadlen MN & Newsome WT (1998) The Variable Discharge of Cortical Neurons: Implications for connectivity, computation, and information coding. The Journal of neuroscience 18(10):3870–3896.

38. van Vreeswijk C & Sompolinsky H (1996) Chaos in neuronal networks with balanced excitatory and inhibitory activity. Science 274(5293):1724–1726.

39. Kanashiro T, Ocker GK, Cohen MR, & Doiron B (2017) Attentional modulation of neuronal variability in circuit models of cortex. Elife 6.

40. Zavitz E, Yu HH, Rowe EG, Rosa MG, & Price NS (2016) Rapid Adaptation Induces Persistent Biases in Population Codes for Visual Motion. J Neurosci 36(16):4579–4590.

41. Gutnisky Da & Dragoi V (2008) Adaptive coding of visual information in neural populations. Nature 452(7184):220–224.

42. Benucci A, Saleem AB, & Carandini M (2013) Adaptation maintains population homeostasis in primary visual cortex. Nat Neurosci 16(6):724–729.

43. Kohn A & Smith MA (2005) Stimulus dependence of neuronal correlation in primary visual cortex of the macaque. The Journal of Neuroscience 25(14):3661–3673.

44. Mitchell JF, Sundberg KA, & Reynolds JH (2009) Spatial attention decorrelates intrinsic activity fluctuations in macaque area V4. Neuron 63(6):879–888.

45. Tolhurst DJ, Movshon JA, & Thompson ID (1981) The Dependence of Response Amplitude and Variance of Cat Visual Cortical Neurones on Stimulus Contrast Exp Brain Res 41:414–419.

46. Dean A (1981) the relationship between response amplitude and contrast for cat striate cortical neurones. journal of physiology 318:413–427.

47. Webster MA (2015) Visual Adaptation. Annu Rev Vis Sci 1:547–567.

48. Reynolds JH & Chelazzi L (2004) Attentional modulation of visual processing. Annual Review of Neuroscience 27:611–647.

49. Maunsell JHR (2015) Neuronal Mechanisms of Visual Attention. Annu Rev Vis Sci 1:373–391.

50. Desimone R & Duncan J (1995) Neural mechanisms of selective visual attention. Annual review of neuroscience 18:193–222.

51. Pooresmaeili A, Poort J, Thiele A, & Roelfsema PR (2010) Separable codes for attention and luminance contrast in the primary visual cortex. The Journal of Neuroscience 30(38):12701–12711.

52. Thiele A, Pooresmaeili A, Delicato LS, Herrero JL, & Roelfsema PR (2009) Additive effects of attention and stimulus contrast in primary visual cortex. Cereb Cortex 19(12):2970–2981.

53. Anton-Erxleben K, Stephan VM, & Treue S (2009) Attention reshapes center-surround receptive field structure in macaque cortical area MT. Cerebral cortex 19(10):2466–2478.

54. Martinez-Trujillo JC & Treue S (2002) Attentional modulation strength in cortical area MT depends on stimulus contrast. Neuron 35(2):365–370.

55. Lee J & Maunsell JHR (2010) The effect of attention on neuronal responses to high and low contrast stimuli. Journal of neurophysiology 104(2):960–971.

56. Reynolds JH, Pasternak T, & Desimone R (2000) Attention increases sensitivity of V4 neurons. Neuron 26(3):703–714.

57. Kohn A & Movshon JA (2003) Neuronal Adaptation to Visual Motion in Area MT of the Macaque. Neuron 39(4):681–691.

58. Kohn A & Movshon JA (2004) Adaptation changes the direction tuning of macaque MT neurons. Nat Neurosci 7(7):764–772.

59. Ghodrati M, Zavitz E, Rosa MGP, & Price NSC (2019) Contrast and luminance adaptation alter neuronal coding and perception of stimulus orientation. Nat Commun 10(1):941.

60. Cohen MR & Kohn A (2011) Measuring and interpreting neuronal correlations. Nature Neuroscience 14(7):811–819.

61. Harris KD & Thiele A (2011) Cortical state and attention. Nat Rev Neurosci 12(9):509–523.

62. Scholvinck ML, Saleem AB, Benucci A, Harris KD, & Carandini M (2015) Cortical state determines global variability and correlations in visual cortex. J Neurosci 35(1):170–178.

63. Ruff DA & Cohen MR (2014) Attention can either increase or decrease spike count correlations in visual cortex. Nature Neuroscience 17(11):1591–1597.

64. Vinck M, Batista-Brito R, Knoblich U, & Cardin JA (2015) Arousal and locomotion make distinct contributions to cortical activity patterns and visual encoding. Neuron 86(3):740–754.

65. McGinley MJ, David SV, & McCormick DA (2015) Cortical Membrane Potential Signature of Optimal States for Sensory Signal Detection. Neuron 87(1):179–192.

66. Ruff DA & Cohen MR (2014) Global Cognitive Factors Modulate Correlated Response Variability between V4 Neurons. Journal of Neuroscience 34(49):16408–16416.

67. Erisken S, et al. (2014) Effects of locomotion extend throughout the mouse early visual system. Curr Biol 24(24):2899–2907.

68. Cunningham JP & Yu BM (2014) Dimensionality reduction for large-scale neural recordings. Nat Neurosci 17(11):1500–1509.

69. Denfield GH, Ecker AS, Shinn TJ, Bethge M, & Tolias AS (2018) Attentional fluctuations induce shared variability in macaque primary visual cortex. Nat Commun 9(1):2654.

70. Geiger J, Lubke J, Roth A, Frotscher M, & Jonas P (1997) Submillisecond AMPA Receptor-Mediated Signaling at a Principal Neuron–Interneuron Synapse. Neuron 18(6):1009–1023.

71. Xiang Z, Huguenard J, & Prince D (1998) GABAA receptor-mediated currents in interneurons and pyramidal cells of rat visual cortex. the Journal of Physiology 506(3):715–730.

72. Salin P & Prince D (1996) Spontaneous GABAA receptor-mediated inhibitory currents in adult rat somatosensory cortex. the Journal of Neurophysiology 75(4):1573–1588.

73. Angulo M, Rossier J, & Audinat E (1999) Postsynaptic Glutamate Receptors and Integrative Properties of Fast-Spiking Interneurons in the Rat Neocortex. the Journal of Neurophysiology 82(3):1295–1302.

74. Levy RB & Reyes AD (2012) Spatial profile of excitatory and inhibitory synaptic connectivity in mouse primary auditory cortex. J Neurosci 32(16):5609–5619.

75. Horvat S, et al. (2016) Spatial Embedding and Wiring Cost Constrain the Functional Layout of the Cortical Network of Rodents and Primates. PLoS Biol 14(7):e1002512.

76. Marino J, et al. (2005) Invariant computations in local cortical networks with balanced excitation and inhibition. Nat Neurosci 8(2):194–201.

77. Sato TK, Haider B, Hausser M, & Carandini M (2016) An excitatory basis for divisive normalization in visual cortex. Nat Neurosci 19(4):568–570.

78. Olsen SR, Bhandawat V, & Wilson RI (2010) Divisive normalization in olfactory population codes. Neuron 66(2):287–299.

79. Verhoef BE & Maunsell JHR (2017) Attention-related changes in correlated neuronal activity arise from normalization mechanisms. Nat Neurosci 20(7):969–977.

80. Ruff DA & Cohen MR (2016) Stimulus Dependence of Correlated Variability across Cortical Areas. Journal of Neuroscience 36(28):7546–7556.

81. Martinez-Trujillo JC & Treue S (2004) Feature-based attention increases the selectivity of population responses in primate visual cortex. Current Biology 14(9):744–751.

82. Maunsell JHR & Treue S (2006) Feature-based attention in visual cortex. Trends in Neurosciences 29(6):317–322.

83. Ruff DA & Born RT (2015) Feature attention for binocular disparity in primate area MT depends on tuning strength. Journal of Neurophysiology 113(5):1545–1555.

84. Treue S & Martinez-Trujillo JC (1999) Feature-based attention influences motion processing gain in macaque visual cortex. Nature 399(6736):575–579.

85. McAdams CJ & Maunsell JHR (2000) Attention to both space and feature modulates neuronal responses in macaque area V4. Journal of neurophysiology 83(3):1751–1755.

86. Moreno-Bote R, et al. (2014) Information-limiting correlations. Nature Neuroscience 17(10):1410–1417.

87. Kanitscheider I, Coen-Cagli R, & Pouget A (2015) Origin of information-limiting noise correlations. Proceedings of the National Academy of Sciences of the United States of America 112(50):E6973–6982.

88. Kohn A, Coen-cagli R, Kanitscheider I, & Pouget A (2016) Correlations and Neuronal Population Information. Annual Review of Neuroscience 39:237–256.

89. Ruff DA, Ni AM, & Cohen MR (2018) Cognition as a Window into Neuronal Population Space. Annu Rev Neurosci 41:77–97.

90. Fries P (2015) Rhythms for Cognition: Communication through Coherence. Neuron 88(1):220–235.

91. Womelsdorf T & Fries P (2007) The role of neuronal synchronization in selective attention. Current opinion in neurobiology 17(2):154–160.

92. Miller EK & Buschman TJ (2013) Cortical circuits for the control of attention. Curr Opin Neurobiol 23(2):216–222.

93. Ni AM, Ruff DA, Alberts JJ, Symmonds J, & Cohen MR (2018) Learning and attention reveal a general relationship between population activity and behavior. Science 359(6374):463–465.

94. Schmolesky M, et al. (1998) Signal timing across the macaque visual system. journal of neurophysiology 79(6):3272–3278.

95. Yu BM, et al. (2009) Gaussian-process factor analysis for low-dimensional single-trial analysis of neural population activity. J Neurophysiol 102(1):614–635.

